# A combined use of intravoxel incoherent motion MRI parameters can differentiate early stage hepatitis-b fibrotic livers from healthy livers

**DOI:** 10.1101/138958

**Authors:** Yì Xiáng J. Wáng, Min Deng, Yáo T. Li, Hua Huang, Jason Chi Shun Leung, Weitian Chen, Pu-Xuan Lu

## Abstract

This study investigated a combined use of IVIM parameters Dslow (*D*), PF (*f*) and Dfast (*D*\*) for liver fibrosis evaluation. 16 healthy volunteers (F0) and 33 hepatitis-b patients (stage F1= 15, stage F2-4 = 18) were included. With a 1.5-T MR scanner and respiration-gating, IVIM diffusion weighted imaging was acquired using a single-shot echo-planar imaging sequence with ten *b*-values of 10, 20, 40, 60, 80, 100, 150, 200, 400, and 800 s/mm^2^. Signal measurement was performed on right liver parenchyma. With a 3-dimensional tool, Dslow, PF, and Dfast values were placed along the x-axis, y-axis, and z-axis, and a plane was defined to separate healthy volunteers from patients. 3-dimensional tool demonstrated healthy volunteers and all patients with liver fibrosis could be separated. Classification and Regression Tree showed a combination of PF (PF < 12.55%), Dslow (Dslow < 1.152 ×10^−3^ mm^2^/s) and Dfast (Dfast <13.36 ×10^−3^ mm^2^/s) could differentiate healthy subjects and all fibrotic livers (F1-F4) with an area under the curve of logistic regression (AUC) of 0.986. The AUC for differentiation of healthy livers vs. F2-4 livers was 1. PF offered the best diagnostic value, followed by Dslow; however, all three parameters of PF, Dslow, and Dfast contributed to liver fibrosis detection.

---

Chronic liver disease is a major public health problem worldwide. The epidemic trend of chronic liver disease is expected to increase owing to an aging population, the growing epidemic of obesity and non-alcoholic steatohepatitis (NASH). Viral hepatitis is the most common blood-borne infection worldwide.^1, 2^ Chronic viral hepatitis can lead to hepatic fibrosis, cirrhosis and hepatocellular carcinoma.^3^ Liver fibrosis, a common feature of almost all chronic liver diseases, involves the accumulation of collagen, proteoglycans, and other macromolecules in the extracellular matrix.^4^ Clinically liver fibrosis usually has an insidious onset and progresses slowly over decades. Originally considered to be irreversible, hepatic fibrosis is now regarded as a dynamic process with the potential for regression.^4^ Treatment with combined therapies on underline etiology and fibrosis simultaneously might expedite the regression of liver fibrosis and promote liver regeneration.^5–7^ Earlier stage liver fibrosis is more amenable to therapeutic intervention. Even when the underline etiology of liver fibrosis could not be eradicated, therapies on liver fibrosis might help delay the progression of the disease to cirrhosis.

To date, noninvasive diagnostic tests available from clinical practice are not sensitive or specific enough to detect occult liver injury at early stage.^8^ Liver biopsy is currently the standard of reference for the diagnosis and staging of liver fibrosis. However, liver biopsy is an invasive procedure with several contraindications and with a risk of complications such as pain, hemorrhage, bile peritonitis, penetration of abdominal viscera, pneumothorax and even death.^9, 10^ The mortality rate associated with needle biopsy was estimated to be between 0.009% and 0.12%.^10^ A noninvasive and quantitative technique for detecting liver fibrosis is highly desirable.

In diffusion-weighted (DW) MRI, the intensity of the acquired magnetic resonance signal depends on the self-diffusion of the excited spins, i.e., on the microscopic stochastic Brownian molecular motion, and the extent and orientation of molecular motion is influenced by the microscopic structure and organization of biological tissues.^11–14^ Perfusion can contribute to the diffusion measurements significantly because of the incoherent motion of blood in pseudorandom capillary network at the macroscopic level.^15–18^ Intravoxel incoherent motion (IVIM) reflects the random microscopic motion that occurs in voxels on MR images of water molecules (either intra-cellular or extracellular) and the microcirculation of blood. In 1986, Le Bihan *et al*^15, 16^ proposed the principle of IVIM which enables the quantitative parameters that separately reflect tissue diffusivity and tissue microcapillary perfusion to be estimated. IVIM signal attenuation is modeled according to the equation

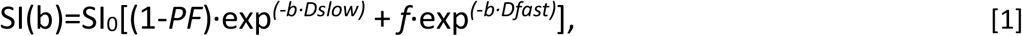

where SI(b) and SI_0_ denote the signal intensity acquired with the *b*-factor value of *b* and *b*=0 s/mm^2^, respectively. Perfusion fraction (PF, or *f*) is the fraction of the pseudo-diffusion linked to microcirculation, Dslow (or *D*) is the true diffusion coefficient representing the pure molecular diffusion (slow component of diffusion), and Dfast (*D*\*) is the pseudo-diffusion coefficient representing the incoherent microcirculation within the voxel (perfusion-related diffusion, or fast component of diffusion).

Molecular water diffusion in fibrotic liver would be restricted by the presence of collagen fibers in the distorted lobular structure. Given the relatively high blood volume fraction of <25–30 mL of blood per 100g in liver,^19^ perfusion can contribute to the diffusion measurements significantly because of the incoherent motion of blood in pseudorandom capillary network at the macroscopic level. It is well accepted that liver fibrosis is associated with reduced liver perfusion.^20–23^ Recently there has been greater interest of using IVIM technique to study diffused liver diseases such as liver fibrosis. However, so far the literatures showed IVIM was unable to detect early liver fibrosis reliably.^24^ We noticed that the potential optimal combination of three IVIM parameters, i.e. Dslow, PF and Dfast, for the detection of liver fibrosis has not been explored in sufficient details. In this study, we set out to explore whether a combination of Dslow, PF and Dfast can be used to separate fibrotic livers from healthy livers. We re-analyze our previously reported cohort, *the Shenzhen 2012/2103 ivim dataset*,^25^ using our updated understanding for IVIM technique and liver imaging. Our literature review showed the *Shenzhen 2012/2103 ivim dataset* remained one of the largest datasets ever reported with all the patients had biopsy histopathology grading [Fig 10 of reference 24].

## Material and Methods

The characteristics of the *Shenzhen 2012/2013 dataset* has been previously reported.^25^ The MRI data was acquired during the period from Aug 1, 2012 to Aug 15, 2013. The study was approved by the ethical committee of Shenzhen No. 3 Hospital, and the informed consent was obtained which included secondary analysis for the data acquired. The IVIM images of one volunteer and one patient were adjudged to contain substantial motion artefacts and therefore excluded for analysis. Sixteen healthy volunteers (10 males, 6 females, mean age: 36.4-yrs old; range: 21–79 yrs old) and 33 consecutively viral hepatitis-b patients were included in the current study. The patient cohort had 15 stage F1 subjects (mean age: 31.8 yrs, 22-53 yrs) and 18 stage F2-4 subjects (mean age: 42 yrs, range: 22-53 yrs). The histology diagnosis for liver fibrosis was based on the consensus of the *2000 Xi'an consensus of the Chinese Society of Infectious Disease and Parasitology and the Chinese Society of Hepatology*,^26^ and being very similar to METAVIR score.^27^ Stage 1 (F1) of liver fibrosis is mild fibrosis only seen at the portal area; stage 2 (F2) indicates fibrosis extending out from the portal areas with rare bridges between portal areas, but without the destruction of the lobular structure; stage 3 (F3) of liver fibrosis is severe fibrosis, there is fibrotic bridging between portal areas and between portal areas and center veins; in stage 4 (F4) there are pseudo-lobules formed and this stage is the final stage of cirrhosis. F0 and F1 livers are commonly referred to as without significant hepatic fibrosis; hepatic fibrosis (F2, F3, and F4) are commonly referred to as significant hepatic fibrosis, and F4 is also referred as cirrhosis.^28^ Hepatic fibrosis is considered clinically significant if defined as F2 or greater than F2, and requiring medical attention.^28, 29^

MR imaging was performed with a 1.5-T magnet (Achieva, Philips Healthcare, Best, Netherlands). The IVIM DW imaging sequence was based on a single-shot DW spin-echo type echo-planar imaging sequence, with ten b-values of 10, 20, 40, 60, 80, 100, 150, 200, 400, 800 s/mm^2^ respectively. SPIR technique (Spectral Pre-saturation with Inversion-Recovery) was used for fat suppression. Respiratory-gating was applied in all scan participants and resulted in an average TR of 1500 ms, and the TE was 63 ms. Other parameters included slice thickness = 7 mm, matrix= 124×97, FOV = 375 mm×302 mm, NEX = 2, number of slices = 6. The IVIM signal attenuation was modeled according to the Equation [1]. The same as our last report,^25^ the estimation of Dslow was obtained by a least-squares linear fitting of the logarithmized image intensity at the *b*-values greater than 200 s/mm^2^ to a linear equation. The fitted curve was then extrapolated to obtain an intercept at *b*=0. The ratio between this intercept and the SI_0_, gave an estimate of PF. Finally, the obtained Dslow and PF were substituted into Eq. [1] and were non-linear least-square fitted against all *b*-values to estimate Dfast using the Levenberg-Marquardt algorithm.

All curve fitting algorithms were implemented in an accustom program develop on MatLab (Mathworks, Natick, MA, USA). Regions-of-interest (ROIs) were positioned to cover a large portion of liver parenchyma while avoiding large vessels (Fig 1). For ROI analysis, the IVIM parameters were calculated based on the mean signal intensity of the whole ROI, which has been shown to offer better estimation than pixel-wise fitting when the signal-to-noise of the DW images is low.^30, 31^

**Fig 1.**
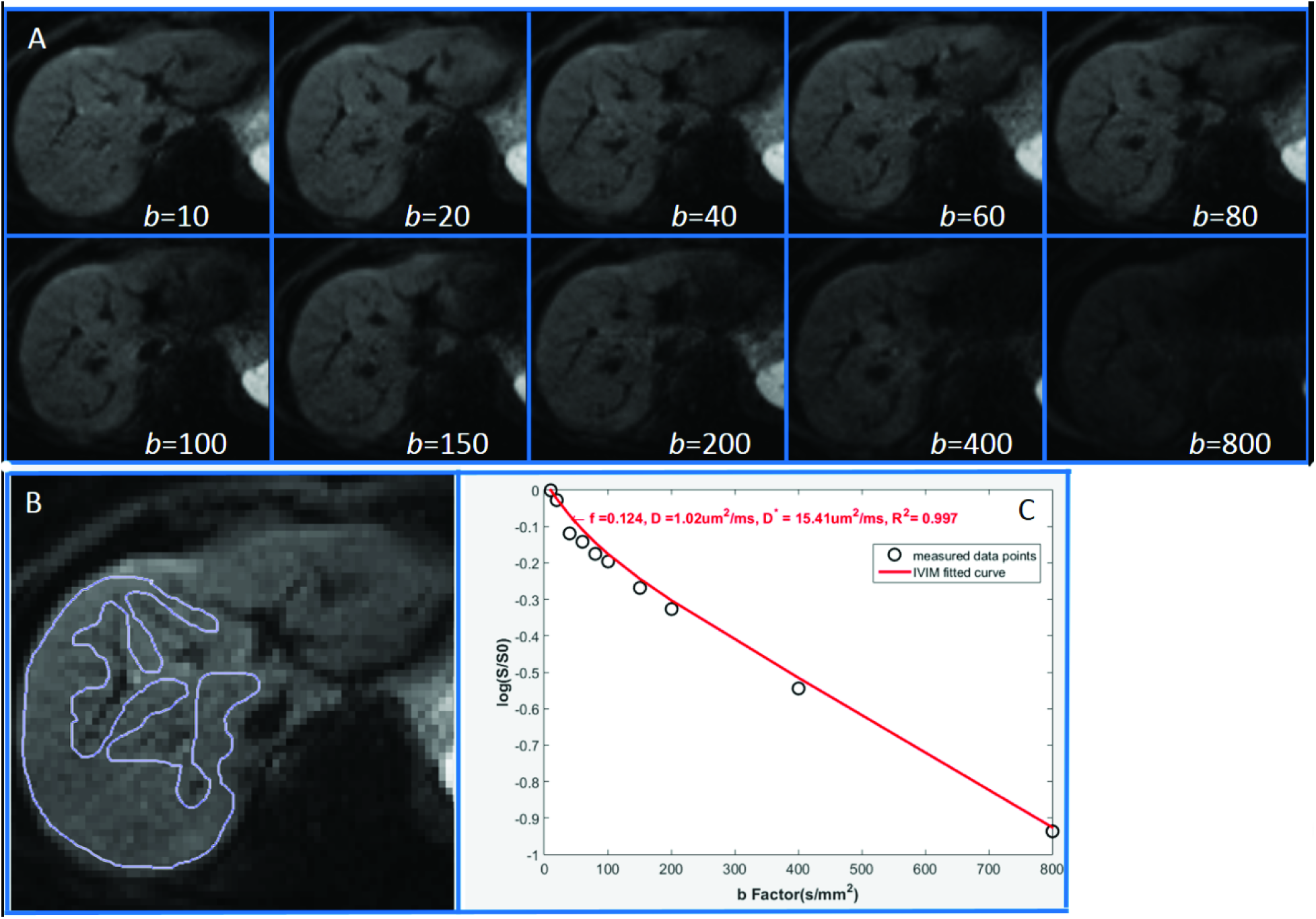
A: Demonstration of a diffusion weighted images with ten b-values from a participant; B: Demonstration of a careful ROI drawing to avoid liver vasculature; C: Signal and *b*-value relationship of the liver slice in B.

The following measurement modifications were made compared with our previous report.^25^ As left lobe of liver is more likely to suffer from artifacts associated the cardiac motion and B0 inhomogeneity susceptibility due its proximity to the stomach and its air inside, therefore in the current study only the right lobe of liver was measured (Fig 1). Figure 1 demonstrates the ROI was carefully drawn to cover only right liver parenchyma while avoiding vasculature and artifacts. All 6 slice per subject were evaluated, while the slices with notable motion artifacts and those demonstrates notable outlier with signal b-value relation were discarded, and finally the slice used for final analysis varied between two to five slices (average: three slices). In addition, with careful histopathology review, two patients with F1 histology score were redefined as into F2. The overall results of current analysis did not differ very notably with previous analysis.^25^

A three-dimensional tool was programed using IBM SPSS 23 for Windows (SPSS Inc., Chicago, IL, USA), and the measures of Dslow, PF, and Dfast were placed along the x-axis, y-axis, and z-axis. Data points from healthy volunteer were labeled as blue in the 3-dimensional space, F1 patients labeled as pink, and F2-4 patients labeled as red. Attempts were then made to separate healthy volunteers from all patients (F1-F4); healthy volunteers from significant patients (F2-F4); and separate patients with different stages. The Support Vector Machine (SVM) approach was used to quantitatively separate the F0 from F1-F4, or F0 from F2-F4.^32^ SVM was used to find a plane (parametrized as Ax + By + Cz + D = 0) that was able to separate the data points into two groups. The distance of the closest data point from an individual group to the separating plane was defined as *d_i_*, where *i* represents the index of the group. The SVM algorithm was used to find an optimal plane which maximizes the margin defined as *d*_1_ + *d*_2_. Prior to calculate the distance *d_i_*, the measured Dslow, PF, and Dfast are normalized by the following equation.

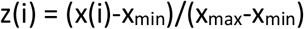

where x(i) is the original data and z(i) is the normalized data; x_max_ and x_min_ are the maximum and the minimum value of x(i), respectively. Note the range of z(i) after normalization is from 0 to 1 for each dimension.

Classification and Regression Tree (CART) model was used to find the cut-off values for PF, Dslow, and Dfast to differentiate F0 livers vs. F1-4 livers and F0 livers vs. F2-4 livers.^33^

## Results

Livers of healthy volunteers had PF of 16.6%±3.6% (mean ± standard deviation), Dslow of 1.14 ± 0.22 (×10^−3^ mm^2^/s), and Dfast of 12.3 ± 3.1 (×10^−3^ mm^2^/s) respectively. The CoV (coefficient of variance, SD/mean) for PF, Dslow and Dfast in healthy volunteers was 0.19, 0.22, and 0.25, respectively.

When the study participants were grouped into three group: 1) healthy volunteers (F0), 2) insignificant liver fibrosis (F1), and 3) significant liver fibrosis (F2, F3, F4), it was seen that PF offered best differentiation of the three group, followed by Dslow (Table 1, Fig-2). By adjusting the viewing angel, the 3-dimensional visual tool demonstrated healthy volunteers (F0, n=16) and all patients with liver fibrosis (F1-4, n=33) could be separated (Fig 3, Fig 4, (supplementary video-1, 2). The cluster of F1 subjects located between F0 and F2-4; however, it was not possible to reliably differentiate patients of different stages.

**Fig 2.**
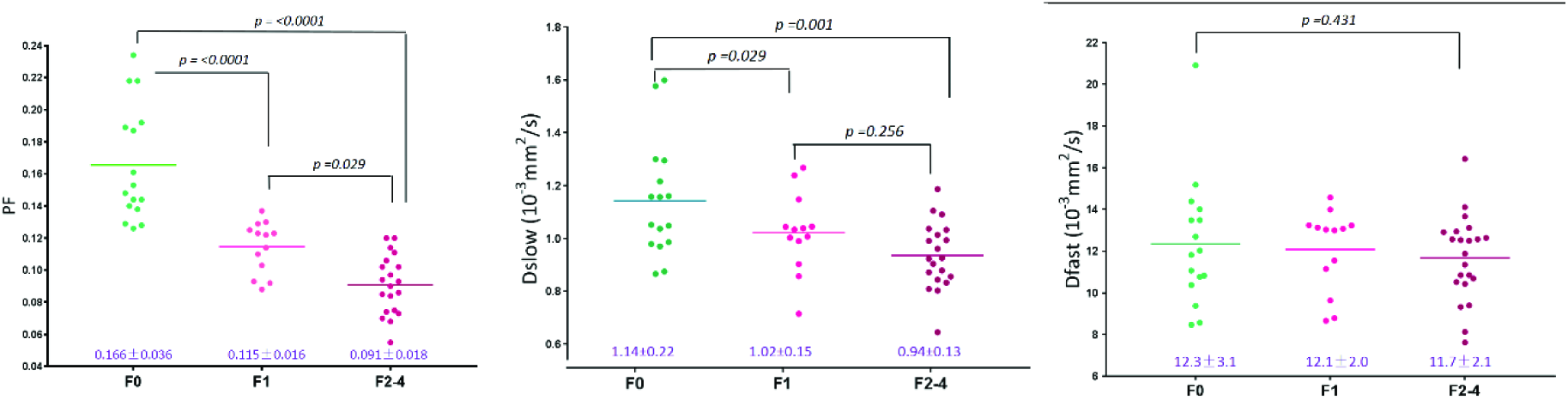
Scattered plots and mean of PF, Dslow, and Dfast of healthy volunteers, F1 liver fibrosis patients, and F2-4 liver fibrosis patients (*p*-value: ANOVA and Mann Whitney U test).

**Fig 3.**
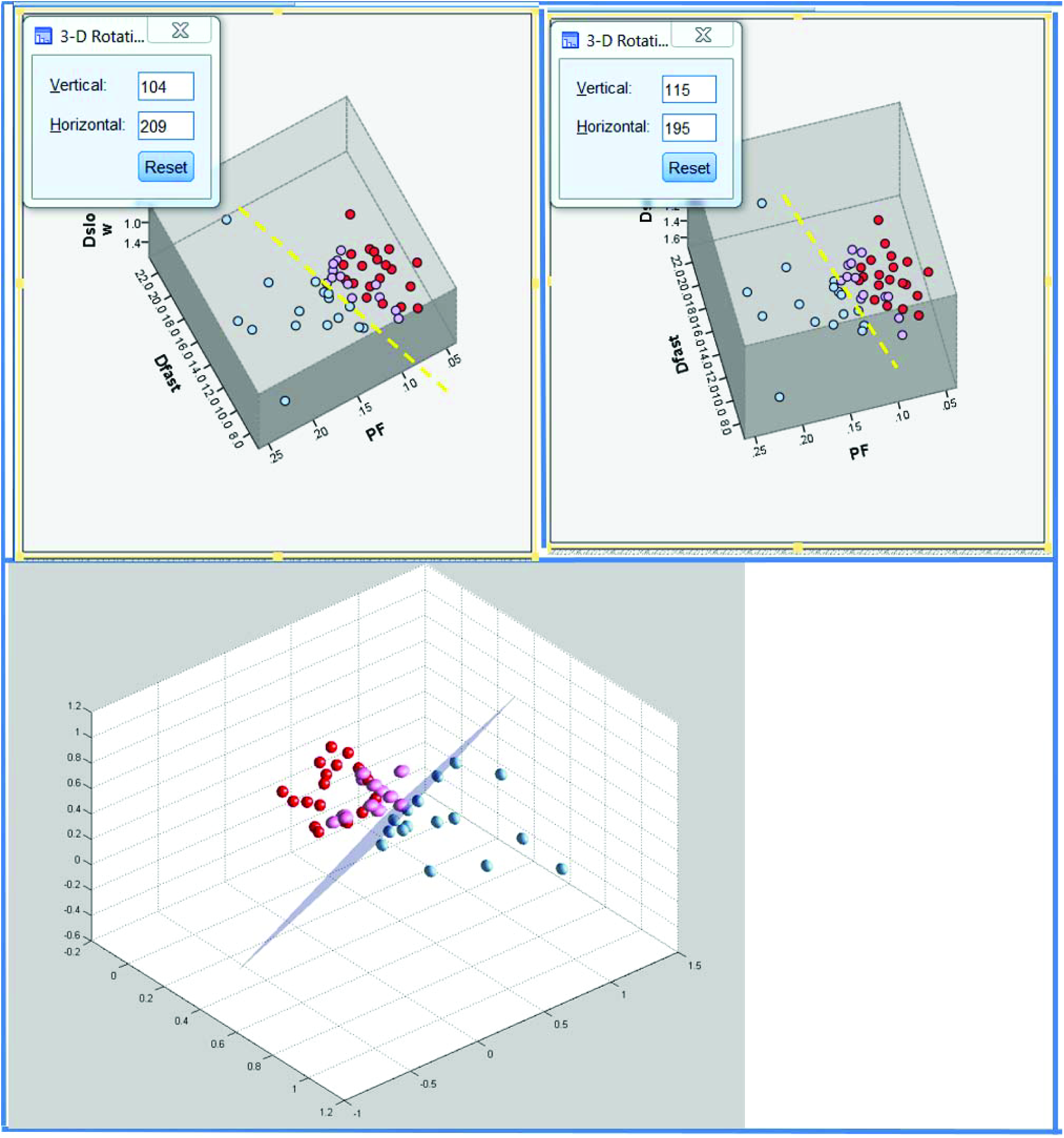
Three-dimensional display of healthy volunteer group (blue balls), F1 patient group (pink balls), and F2-3 patient group (red balls). Each ball represents one participant. The differentiation of volunteer group and patient group can be better visualized by rotating in 3-dimensional space (dotted yellow line or purple plane).

**Fig 4.**
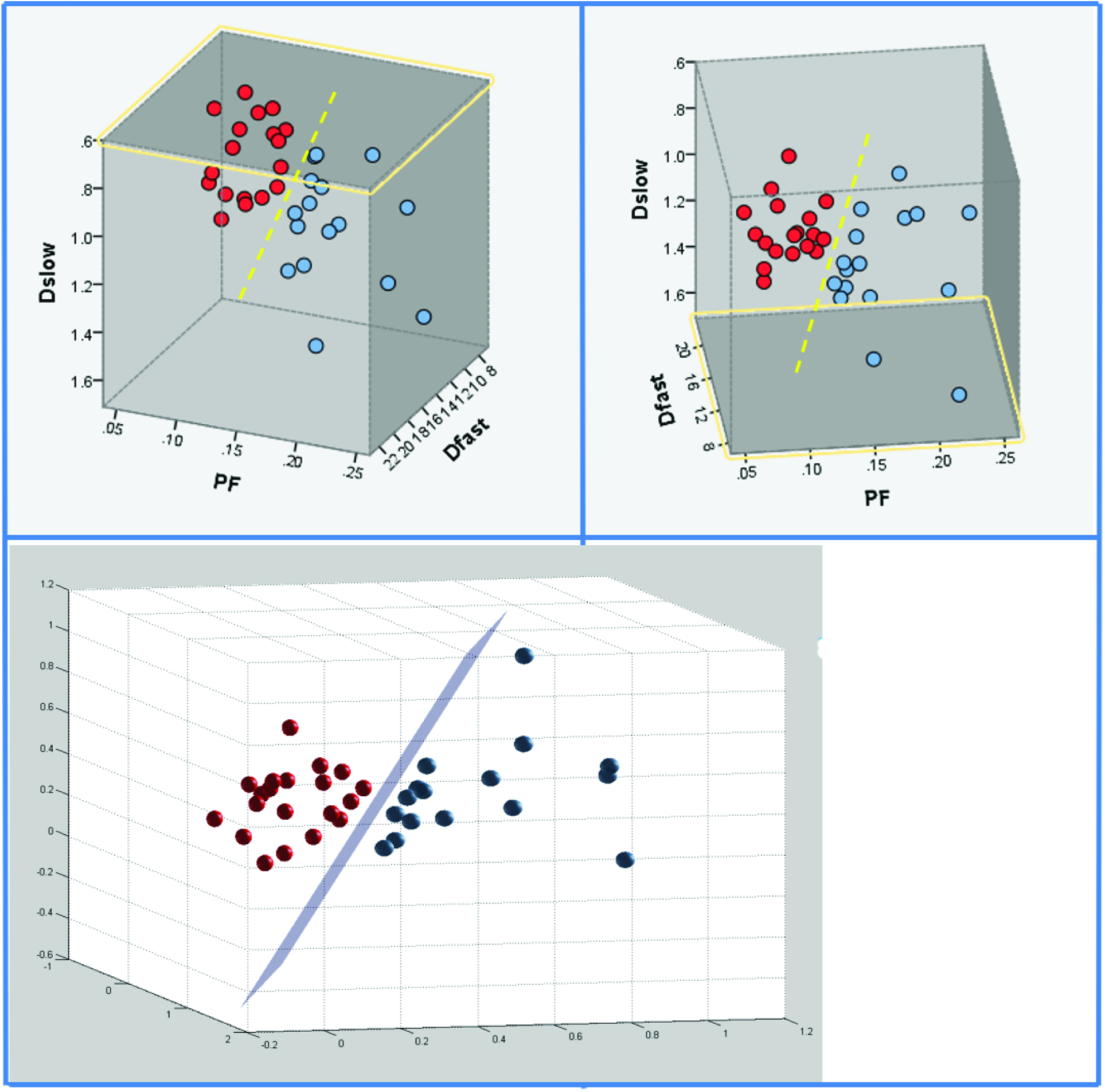
Three-dimensional display of healthy volunteer group (blue balls), and F2-3 patient group (red balls). Each ball represents one participant. The differentiation of volunteer group and patient group can be better visualized by rotating in 3-dimensional space (dotted yellow line or purple plane).

**Table 1.** Mean, standard deviation (sd), and coefficient of variance (CoV) of PF, Dslow, and Dfast of healthy volunteers, F1 liver fibrosis patients, and F2-4 liver fibrosis patients.

| | PF | | | Dslow $\times 10^{-3} \text{ mm}^2/\text{s}$ | | | Dfast $\times 10^{-3} \text{ mm}^2/\text{s}$ | | |
| --- | --- | --- | --- | --- | --- | --- | --- | --- | --- |
|  | F0 | F1 | F2-4 | F0 | F1 | F2-4 | F0 | F1 | F2-4 |
| mean | 0.165 | 0.11 | 0.091 | 1.14 | 1.01 | 0.94 | 12.34 | 12.03 | 11.68 |
| sd | 0.036 | 0.017 | 0.019 | 0.22 | 0.14 | 0.13 | 3.06 | 1.85 | 2.22 |
| CoV | 0.215 | 0.157 | 0.211 | 0.19 | 0.14 | 0.14 | 0.25 | 0.15 | 0.19 |

Quantitative analysis with SVM showed healthy volunteers and all patients with liver fibrosis (F1-4) were differentiated with a plane defined by *(166.58*PF) +(8.90*Dslow) - (0.98*Dfast) - 19.71 = 0* (Fig 3); healthy volunteers and patients with significant liver fibrosis (F2-4) were more clearly differentiated with a plane defined by *(29.56*PF) + (4.33*Dslow) - (0.12*Dfast)-6.67 = 0* (Fig 4). The mean distance of the data points for F0 vs F1-4 to the central plane was *0.0021_F0_+0.0026_F1-2_=0.0047*, and for F0 and F2-4 to the central plane was *0.0149_F0_+0.0138_F2-4_= 0.0287*.

Three-dimensional visual tool demonstrated better differentiation of healthy livers and fibrotic livers than two-dimensional plot using PF and Dslow values (Fig 3–5), indicating Dfast contributed to differentiating healthy volunteers and patients with liver fibrosis.

**Fig 5.**
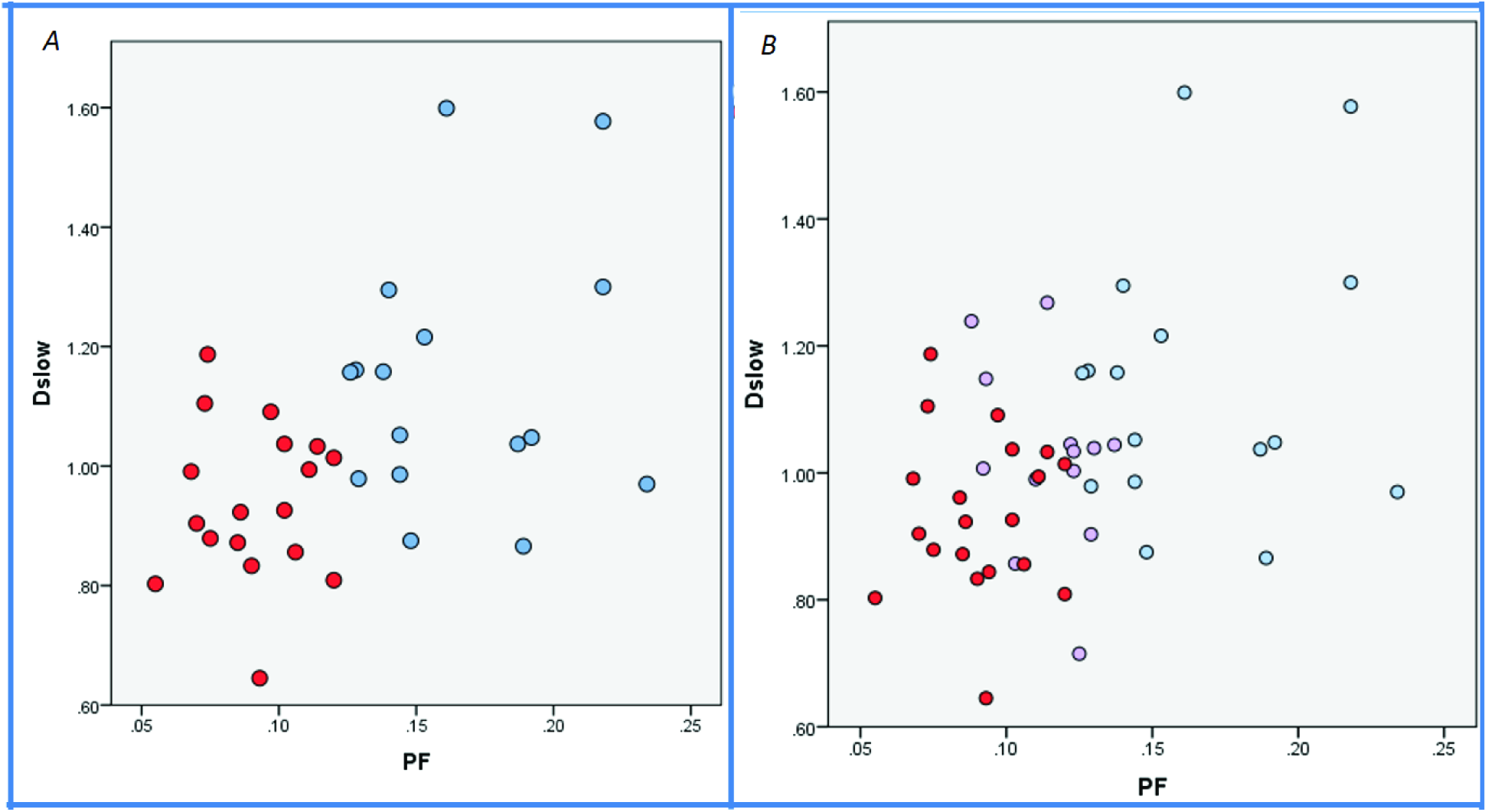
Two-dimensional demonstration of healthy volunteer group (blue balls), F1 patient group (pink balls), and F2-3 patient group (red balls) using PF-axis and Dslow-axis. Each ball represents one participant. A combination of PF-axis and Dslow-axis is insufficient to differentiate F0 subjects vs F1-4 patients, in contrast to demonstrations in Fig 3 and Fig 4.

The CART analysis result is shown in table 2, a combination of PF (cutoff value: PF < 12.55%), Dslow (Dslow < 1.152 ×10^−3^ mm^2^/s) and Dfast (Dfast<13.36 ×10^−3^ mm^2^/s) can differentiate healthy subjects (F0) and fibrotic livers (F1-F4) with an AUC (Area under the curve of logistic regression) of 0.986. The AUC for differentiation of healthy subjects (F0) vs. significantly fibrotic (F2-4) was 1.

**Table 2.** CART analysis of area under the curve of logistic regression (AUC) of F1-4 or F2-4 (comparing with F0) on PF, Dslow and Dfast

| Model | AUC (1): F1-4 vs F0 | AUC (2): F2-4 vs F0 |
| --- | --- | --- |
| PF | 0.9545 (cutoff value: $PF < 0.1255$) | 1 (cutoff value: $PF < 0.123$) |
| PF and Dslow | 0.9773 (cutoff value: $D_{\text{slow}} < 1.152$) | 1 (cutoff value: $D_{\text{slow}} < 1.131$) |
| PF, Dslow and Dfast | 0.9858 (cutoff value: $D_{\text{fast}} < 13.36$) | 1 (cutoff value: $D_{\text{fast}} < 13.3$) |

Unit of Dslow and Dfast: $\times 10^{-3} \text{ mm}^2/\text{s}$

## Discussion

Currently the most clinically used imaging technique for evaluation of liver fibrosis is ultrasound elastography, while the investigational technique of MR elastography has undergone many clinical trials.^34^ US elastography method is inexpensive, fast to acquire, and do not require postprocessing. In particular, 1-dimensional transient elastography has been adopted clinically. The diagnostic accuracy of ultrasound elastography has been assessed in numerous studies and pooled in meta-analyses. The reported diagnostic accuracy for various ultrasound elastography techniques has an AUC in the range of 0.84–0.87 for fibrosis stage ≥ F2, 0.89–0.91 for fibrosis stage ≥ F3, and 0.93–0.96 for fibrosis stage F4.^35–37^ MR Elastography provides higher overall diagnostic accuracy than ultrasound-based elastography.^34^ Meta-analyses report AUC for MR Elastography in the range of 0.84–0.95 for diagnosing fibrosis stage ≥ F1, 0.88–0.98 for fibrosis stage ≥ F2, 0.93–0.98 for fibrosis stage ≥ F3, and 0.92–0.99 for fibrosis stage F4.^38–40^ The limitations of ultrasound elastography include it is an operator-dependent technique; while MR Elastography requires additional hardware (elastography driver) to be added to the MR scanner. As IVIM imaging sequence is widely available in clinical MR scanners, it represents a promising alternative to existing techniques for liver fibrosis evaluation.

The PF and Dslow measurements obtained in this study broadly agreed with previous reports.^24, 41–46^ For the 27 studies which reported measurement for healthy livers, the median value for Dslow was 1.11×10^−3^ mm^2^/s and 1.02×10^−3^ mm^2^/s at 1.5 and 3 T respectively; the median value for PF was 22.00% and 22.65% at 1.5 and 3T respectively.^24^ Using a 25 b-values acquisition, ter Voert *et al*^41^ reported CoV of 0.23, 0.22, and 0.68 for PF, Dslow and Dfast of normal liver regions of 15 subjects. In other five reports, CoV for PF and Dslow in healthy subjects have been reported to be 0.13 and 0.11,^42^ 0.30 and 0.21,^43^ 0.28 and 0.19,^44^ 0.20 and 0.64,^45^ 0.18 and 0.05.^46^ For healthy subjects, the CoV in our study was 0.21, 0.19 and 0.25 for PF, Dslow and Dfast respectively. Dfast value obtained in this study is indeed lower than most of the previous reports.^24^ The computing of Dfast, and to a less extent also the PF, is expected to have been comprised by that we did not obtain *b*=0 images for the *Shezhen201/2013 ivim dataset* (for an example with *b*=0 image included, see supplementary document 1). Dfast is more associated with the low *b*-values (<50 s/mm^2^), which corresponds to the steep part in the measured signal versus b-value relationship (Fig 5 of reference 24).

Though many researches have been published on the evaluation of liver fibrosis using IVIM, how to optimally combine Dslow, PF, and Dfast to get diagnostic information is not yet explored. The most important result of the current study is that despite the *Shezhen201/2013 ivim dataset* was not acquired with an optimized protocol, still we were able to demonstrate that healthy volunteers and patients with liver fibrosis could be differentiated. Another important finding of this study is that among the IVIM parameters of PF, Dslow and Dfast, PF offers best diagnostic value; and Dfast can provide additional differentiation value though it is a less stable measurement. Overall the cluster of F1 subjects located between F0 and F2-4; however, it was not possible to differentiate patients of different stages of F2-F4 in this study. We expect this is at least partially due to the fact that the histological diagnosis is also not a clear-cut, a high end F1 liver will be similar to a low end F2 liver.^47^ The findings of this study are important as till now it has been considered that there is no reliable noninvasive method, being imaging or serum biomarkers, can reliably detect early stage liver fibrosis. In the meantime, we are looking into further validating our approach with other dataset or with new prospective studies. Additionally, a 2-dimensional flat plane was used in this study to separate healthy livers vs. fibrotic livers, theoretically curved planes can also be used if curved planes offer better separation of healthy livers vs. fibrotic livers or better staging of fibrotic livers.

IVIM parameters strongly depend upon the choice of the b-values and the threshold used for computation. Numerical modeling suggests that the estimation uncertainties of Dslow and PF can reach 3.89% and 11.65% respectively with typical parameter values at a moderate signal-to-noise ratio (SNR) of 40. ^48^

However, to estimate Dfast within 10% uncertainty requires SNR > 122 [41]. Pekar *et al*.^49^ commented that Dfast in particular tends to be unstable unless an unrealistically high SNR is achieved. More *b*-values and applying an optimized b-value distribution reduce errors in the IVIM parameter estimation.^24, 40, 50–53^ In study of ter Voert *et al* the imaging time for IVIM with 25 *b*-values was 5 to 6 minutes which is clinically acceptable. Empirical literatures also suggest that Dslow is the most reliable parameter among the three parameters.^24^ However, Dslow may suffer from limited dynamic range for detecting fibrotic changes in the liver as shown this study. On the other hand, PF may offer both reasonable measurement stability and sufficient dynamic range.

There are a few ways to improve the measurement accuracy in this study. This study used only 10 *b*-values, while ter Voert *et al* recommend no less than 16 b-values for IVIM quantification to improve the measurement reliability.^34^ The cut-off *b*-value to obtain Dfast for this study, i.e. *b*=200, might have been too high.^24^ It will be worthwhile to test to assign only *b*-values of less than 50 s/mm^2^ as low b-values. The cutoff value of PF <12.55%, Dslow < 1.152 ×10^−3^ mm^2^/s and Dfast < 13.36 ×10^−3^ mm^2^/s (table 2) may need to be re-adjusted if different b-value distribution is used. DW imaging is very sensitive to any macroscopic patient motion. Due to the extensive residual motion from respiratory gated data acquisition^24^, it may be beneficial to use breath-hold technique even at the cost of reduced b-value number or reduced voxel spatial resolution. Our previous experience suggests that it is possible to get precise liver tissue measurement by multiple breath-hold.^54^ Other approaches will be to de-noise as well as design better segmentation to statistically remove ill-fitted pixels in ROI, and employ better fitting strategies and motion correction.^55–59^ Actually, our preliminary analysis demonstrated IVIM parameters’ scan-rescan reproducibly can be satisfactory if motion contaminated data are carefully removed during analysis. Another limitation of our study is all our patients had liver fibrosis due to viral hepatitis-b. Whether results of our study can be generalized to liver fibrosis of other causes, such as NASH, remains to be validated. Quantification of diffusion may be confounded by fat and iron in the liver,^60^ and this has not been carefully investigated in this study. However, it has been already shown that imaging can reliably detect late stage liver fibrosis and liver cirrhosis,^61^ and the question which requires more researches is how to detect F1 and F2 stage liver fibrosis. Additionally, it is known that liver fat and iron can be quantified by MRI.^62, 63^ It is expected that with better IVIM imaging protocol with more b-values and better image post-procession, differentiation of early stage fibrotic liver from healthy liver should have increased reliability. The use of Bayesian prediction, incorporating relevant findings from the available methods, is also a promising technique for liver fibrosis evaluation.^64^ The Bayesian prediction provides probabilities, rather than a ‘yes/no’ decision; it also allows weighting of the different methods, such as IVIM, liver T1rho^65, 66^, and elastography^38–40^ readouts, therefore realizing multi-parameter diagnosis.

In conclusion, a combination of PF, Dslow and Dfast shows the potential of IVIM to detect early stage liver fibrosis. Among the three parameters PF offers best diagnostic value, followed by Dslow; however, all three parameters contribute to liver fibrosis evaluation. Further researches shall improve image data post-processing, denoise or remove poorly fitted regions in the liver, and validate our approach with additional datasets.

Footnote: *The Shenzhen 2012/2103 ivim dataset* is available to external researchers for analysis upon contacting the corresponding author of this article.

## Acknowledgement

This study was supported by a direct grant for research from the Chinese University of Hong Kong (No. 4054167). The authors thank Dr Jing Yuan, Medical Physics and Research Department, Hong Kong Sanatorium and Hospital, for setting up the data acquisition protocol at the Shenzhen No. 3 People's Hospital.

Supplementary video-1: Three-dimensional space video rendering of the relationship among normal liver (blue balls), early stage liver fibrosis (F1, pink balls), and significant liver fibrosis (F2-4, red balls). It is possible to differentiate healthy volunteers and all patients with liver fibrosis.

Supplementary video-2: Three-dimensional space video rendering of the relationship between normal liver (blue balls) and significant liver fibrosis (F2-4). Healthy volunteers and patients with significant liver fibrosis can be reliable differentiated.

